# *E. coli* adhesion and biofilm formation on polydimethylsiloxane are independent of substrate stiffness

**DOI:** 10.1101/2020.01.15.907956

**Authors:** Sandra L. Arias, Joshua Devorkin, Ana Civantos, Jean Paul Allain

## Abstract

Bacterial adhesion and biofilm formation on the surface of biomedical devices is a detrimental process that compromises patient safety and material functionality. Several physicochemical factors are involved in biofilm growth, including the surface properties. Among those, material stiffness has recently been suggested to influence microbial adhesion and biofilm growth in a variety of polymers and hydrogels. However, no clear consensus exists about the role of material stiffness on biofilm initiation and whether very compliant substrates are deleterious to bacterial cell adhesion. Here, by systematically tuning substrate topography and stiffness while keeping the surface free energy of polydimethylsiloxane substrates constant, we show that topographical patterns at the micron and submicron scale impart unique properties to the surface that are independent of the material stiffness. The current work provides a better understanding of the role of material stiffness on bacterial physiology and may constitute a cost-effective and simple strategy to reduce bacterial attachment and biofilm growth even in very compliant and hydrophobic polymers.

## Introduction

Infections associated with the use of implantable and prosthetic devices are responsible for about half of all nosocomial infections in the United States, and despite continued efforts in the field, device-associated infections are still the most prevalent cause of biomaterial failure^1^. Because of their abiotic and avascular nature, implantable and prosthetic devices can be quickly colonized by microorganisms that usually inhabit epithelial tissues and mucous membranes like the skin, periurethral area, and mouth^1,2^. Indeed, abiotic surfaces are colonized by 10,000 times less bacterial load than that needed to invade native tissues^1,3^. Once pathogenic microorganisms colonize the surface of biomaterials, they aggregate into microbial communities enclosed by extracellular polymeric substances known as a biofilm. Biofilms increase the resistance of microorganisms to antibiotics and lead to life-threatening outcomes such as the dissemination of bacteria to other body sites, chronic inflammation, and bacteremia^3,4^. In intensive care units, for example, bacterial contamination of intravenous catheters and indwelling urethral catheters costs about $300 million to $2.3 billion annually. Those expenses can scale quickly as a result of the patient’s extended hospital stays and associated treatment^4,5^.

Bacterial adhesion on abiotic surfaces is the first step in biofilm initiation; thus, elucidating the events that influence adhesion is pivotal to design strategies that can prevent biofilm-associated infections. Bacterial adhesion is a complex process known to depend on several physicochemical factors, including the biological properties of the microorganism and the material interface itself ^6,7^. Bacteria adhesion mechanisms have been associated with various adhesins and organelles (e.g., fimbriae, flagella). Moreover, protein complexes located at the outer membrane can also adhere directly to abiotic surfaces without the need of a pre-conditioned protein layer^6,8^. The material physicochemical properties that have been shown to influence bacterial adhesion include surface chemistry, surface charge density, surface energy, and more recently, the topographical features and the mechanical properties of the material^7,9^. The effect of surface topography on bacterial adhesion has been demonstrated by mimicking the skin of marine animals that resist high levels of biofouling in seawaters, including that of the starfish and shark’s skin. For example, several authors have found a reduction in microbial adhesion and settlement, swarming motility, and biofilm development when using surface microtopographies with feature size comparable or slightly larger than that of the fouling microorganisms^10–13^.

The mechanisms behind the anti-biofouling capabilities of surface microtopographies are not fully understood. In general, topographical patterns have been proven to affect the settlement preferences in a variety of microorganisms, and influence its adhesion strength, attachment density, and distribution^14^. It has been suggested that microorganisms tend to maximize the contact area with a surface^15^, or preferentially bind to locations that maximize the number of attachment points^16^. Recently, these mechanisms of selectivity have been associated with a cost of cell adhesion, since surface arrangements containing local curvature at the micron and submicron scale (e.g., hemispheres patterns) seem to be unfavorable for bacterial attachment despite their large surface area^11,17^. Moreover, using recessed patterns inspired by the Shark’s skin, Sakamoto and coworkers^18^ showed that the geometric complexity of the surface (e.g., tortuosity) rather than the feature depth in grooved patterns, could reduce biofilm formation and the swarming motility in both *Pseudomonas aeruginosa* and *Staphylococcus aureus* strains.

Like topographical features, the surface stiffness has also recently been suggested to impact bacterial adhesion. However, contradictory results have been obtained for a range of synthetic polymers and hydrogels, and it is still debatable whether material stiffness influences bacterial adhesion and subsequent biofilm formation. For example, by varying the monomer to crosslinker ratio, Kolewe and coworkers^19^ studied the attachment density of *Escherichia coli* and *Staphylococcus aureus* as a function of Young’s modulus in poly(ethylene glycol)dimethacrylate and agar. These authors found that fewer bacteria attached to softer hydrogels and suggested that material stiffness could potentially reduce the initial adhesion of bacteria on both synthetic polymers and natural hydrogels. In sharp contrast, Wang and coworkers^20^ found that soft substrates had the highest initial rate of *S. aureus* deposition on polyacrylamide (PAAm) hydrogels. In opposition to the above observations, Straub and coworkers^21^ have proposed that the viscosity and hydrophobic interactions, not the material stiffness, are responsible for the differences in bacterial adhesion reported on those polymers.

In this study, we aimed to shed light on these disparities by examining the role of substrate stiffness on the adhesion and biofilm formation of *E. coli* to poly(dimethylsiloxane) (PDMS). PDMS is a crosslinked polymer commonly used in a range of biomedical applications, including medical adhesives, catheters, blood pumps, mammary prosthesis, drug delivery systems, among many others^22^. We used low-energy singly-charged inert ions to irradiate the PDMS and yield substrates with variable stiffness but comparable surface free energy. This process also resulted in the formation of a wavy-like topography (or wrinkles) in the PDMS, whose dimensions were controlled by the irradiation parameters. We then examined the influence of material stiffness on adhesion and biofilm formation of *E. coli*, independent of the wetting properties and surface free energy of the substrates. By systematically varying the wrinkle size and stiffness of PDMS, we showed that complex submicron topographies limited the adhesion of *E. coli* to PDMS independent of the substrate stiffness. We also showed that for stiff and very compliant PDMS, cell adhesion and biofilm mass were reduced by controlling the surface topography at the micron and submicron scale. Our findings contribute to a better understanding of the role of material stiffness on cell adhesion and biofilm formation. Because *E. coli* is a significant source of medical-device acquired infections, our results provide a cost-effective and simple way to reduce bacterial attachment before biofilms can be formed, even in very compliant and hydrophobic polymers.

## Materials and Methods

### Preparation of PDMS substrates

Microscope glass slides were cut in squares of 15 mm side and cleaned in a solution of 1:3 of hydrogen peroxide to sulfuric acid for 1 h. Glass slides were subsequently rinsed with milli-Q water several times. Excess of water was removed by flowing the glass with nitrogen gas just before spin coating. Polydimethylsiloxane (PDMS) (Dow SYLGARD™ 184 silicone elastomer) was prepared by mixing the liquid silicone pre-polymer and the curing agent at ratios of 10:1, 30:1 and 50:1. Bubbles formed in the PDMS were removed by placing the preparations in a desiccator connected to the vacuum line for 15 min. A spin coater (Specialty Coating Systems Model 6808) operating at 1500 and 2500 rpm were used to deposit the PDMS on the cleaned glass slides. PDMS was then cured at 80°C for 45 min and used the next day for nanopatterning.

### Argon irradiation of polydimethylsiloxane

Spin-coated PDMS films deposited on glass sliders were inserted in a vacuum chamber and irradiated with argon (Ar^+^) ions. The fixed parameters during the irradiation were: ion species (Ar^+^), ion energy (1 KeV), and fluence (1×10^17^ ions/cm^2^). A series of experiments varying the angle of irradiation (0°, 45°, and 75°) and the stiffness of the PDMS substrates (≈ 0.01, 0.1, and 1 MPa) were performed to yield wrinkles with variable amplitude and wavelength. The irradiations were performed using an ion current in the range of 0.3 to 0.6 mA. The base pressure of the vacuum chamber was 2−10^−6^Pa.

### Atomic force microscopy (AFM)

An Asylum Cypher AFM was used for the topographical characterization of the argon-irradiated PDMS. The instrument was operated in tapping mode in air using a BS-Tap300AI cantilever (BudgetSensors) at scan sizes of 5, 10, and 20 μm. The surface roughness power spectral density was obtained by computing the radially averaged two-dimensional power spectrum of the topography using a MATLAB routine developed by Mona Mahboob Kanafi and available on the MathWorks website page^23^.

### Nanoindentation

The Optics11 Piuma soft material indenter was used to determine the Young modulus of the PDMS before and after irradiation. The stiffness of PDMS substrates prepared at a ratio of 10: 1 was determined with a spherical glass bead of 8 μm in radius and a cantilever spring constant of 46.900 N/m. Soft PDMS typical of preparations at ratios of 30:1 and 50:1 were analyzed in ultrapure water to reduce the bead adhesion to the substrate during indentation. The probe used on these samples had a spherical glass bead of 9.5 μm in radius and cantilever spring constant of 4.000 N/m. At least 25 indentations were performed per condition, using a step size of 25 μm between indentation points.

### Water contact angle and surface free energy

The wetting properties of untreated and wrinkled PDMS were measured via static contact angle within 48 h after argon irradiation using a Rame-Hart Model 250 Standard Goniometer. Ultrapure water droplets of 2 to 4 μl in volume were used for the water contact angle measurements using a precision syringe. A total of 5 to 6 droplets were placed per sample and analyzed using the instrument’s software. For the surface free energy determination, droplets of diiodomethane were created and analyzed for each experimental sample following the same procedure as with the water droplets. The surface free energy was then computed by solving the Owens and Wendt^24^ equation:

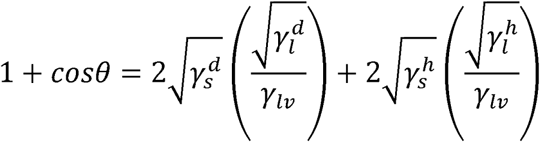

Where *θ* is the contact angle of a liquid on a solid, *γ*_*lv*_ is the free energy of the liquid against its saturated vapor (*v*), and the superscripts *d* and *h* refer to the hydrogen bonding and dispersion force components of the free energy of the liquid. The subscripts *s* and *l* denote the solid and liquid, respectively. By measuring the contact angle of

diiodomethane and water and using known values of 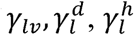; the equation can be solved for 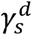 and 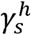.

### Bacterial strains and culture conditions

*Escherichia coli* constitutively expressing the Green Fluorescent protein (ATCC^®^ 25922 GFP) was grown overnight at 37°C with shaking in a tryptic soy broth medium containing ampicillin at an optical density of 0.5-0.6. Before each experiment, the overnight culture was diluted at a 1:10 ratio in an M63 minimal media and incubated in the presence of the untreated and freshly irradiated PDMS samples for 24 h at 30°C without shaking. The M63 minimal media was prepared by diluting 15.6 g of M63 broth powder (VWR Life Science) in 0.9 L ultrapure water, and the pH was adjusted to 7.0 using potassium hydroxide. The solution was sterilized by autoclaving, and 20% sterile glycerol and 1M sterile-filtered MgSO_4_ were added to the media subsequently.

### Crystal Violet Staining

Biofilm mass accumulation formed on untreated and wrinkled PDMS was determined by Crystal Violet staining. For that end, *E. coli* was incubated in the presence of the unmodified and wrinkled PDMS for 24 h at 30°C without shaking, as stated previously. After that, substrates were gently rinsed with phosphate buffer saline three times to remove loosely attached cells and dried for 20 min at room temperature. Subsequently, the experimental samples were stained for 5 min with a solution of 0.1% Crystal Violet prepared in ultra-pure water. Subsequently, the colorant was discarded, and the samples were rinsed several times with ultra-pure water and dry at room temperature for one hour. The stained biofilms were then solubilized with 100% ethanol for 5 min, and the absorbance was read at 570 nm in a 96-well plate using a microplate reader (Biotek Synergy). Each experiment was performed with triplicate samples per condition.

### Laser scanning confocal microscopy

*E. coli* constitutively expressing the Green Fluorescent protein present on biofilms was examined by laser scanning confocal microscopy (Leica TCS SP8) using oil immersion at 40X magnification. To determine the distribution of the bacterial cells and biofilms respect to the topography, *E. coli* was fixed with 10% formalin and serially dehydrated in 20, 40, 60, 80, and 100% ethanol dilutions in ultra-pure water for 10 min each. Optical images of *E. coli* were obtained using a Keyence VK-X1000 3D laser scanning confocal microscope. Fluorescent and optical Images were captured at several random locations across each sample.

### Statistics

One -way ANOVA and Tukey’s multiple comparisons post-test analysis were applied to determine statistical differences between samples. The significance was set at 5%. Reported values in figures and tables denote the mean ± standard deviation (SD).

## Results

Wrinkles were fabricated with variable amplitude and wavelength on PDMS by tuning the material stiffness and angle of incidence during low-energy ion beam irradiation (**Figure 1**). We adjusted the stiffness of the PDMS by using mixtures of the base to crosslinker of 10:1, 30:1, and 50:1 w/w, to yield substrates with Young’s modulus of about 1, 0.1 and 0.01 MPa, respectively. Notice that the lower the ratio of base to crosslinker (e.g., 10:1), the stiffer the resulting PDMS, and the shorter the period of the wrinkled pattern that formed after low-energy ion beam irradiation (**Figure 1 upper panel**). The wrinkle size was further tuned by increasing the angle of irradiation from a normal incidence (0°) to an oblique angle of 45° while keeping constant the other irradiation parameters (e.g., argon ions (Ar^+^) at 1 keV and fluence of 1×10^17^ cm^-2^) (**Figure 1 lower panel**). The formation of wrinkled patterns in PDMS is known to be a surface instability phenomenon that arises when the mechanical properties of the top layer divergent from those of the bulk material^25^. During ion beam irradiation, the top layer of the PDMS is chemically modified, forming a silica-like thin film that is stiffer than the supporting polymer underneath. Once the ion beam is removed, the stiff top causes the contraction of the unmodified PDMS, giving rise to a wrinkled pattern whose wavelength and amplitude depend on the degree of cross-linking of the top layer^26^.

**Figure 1.**
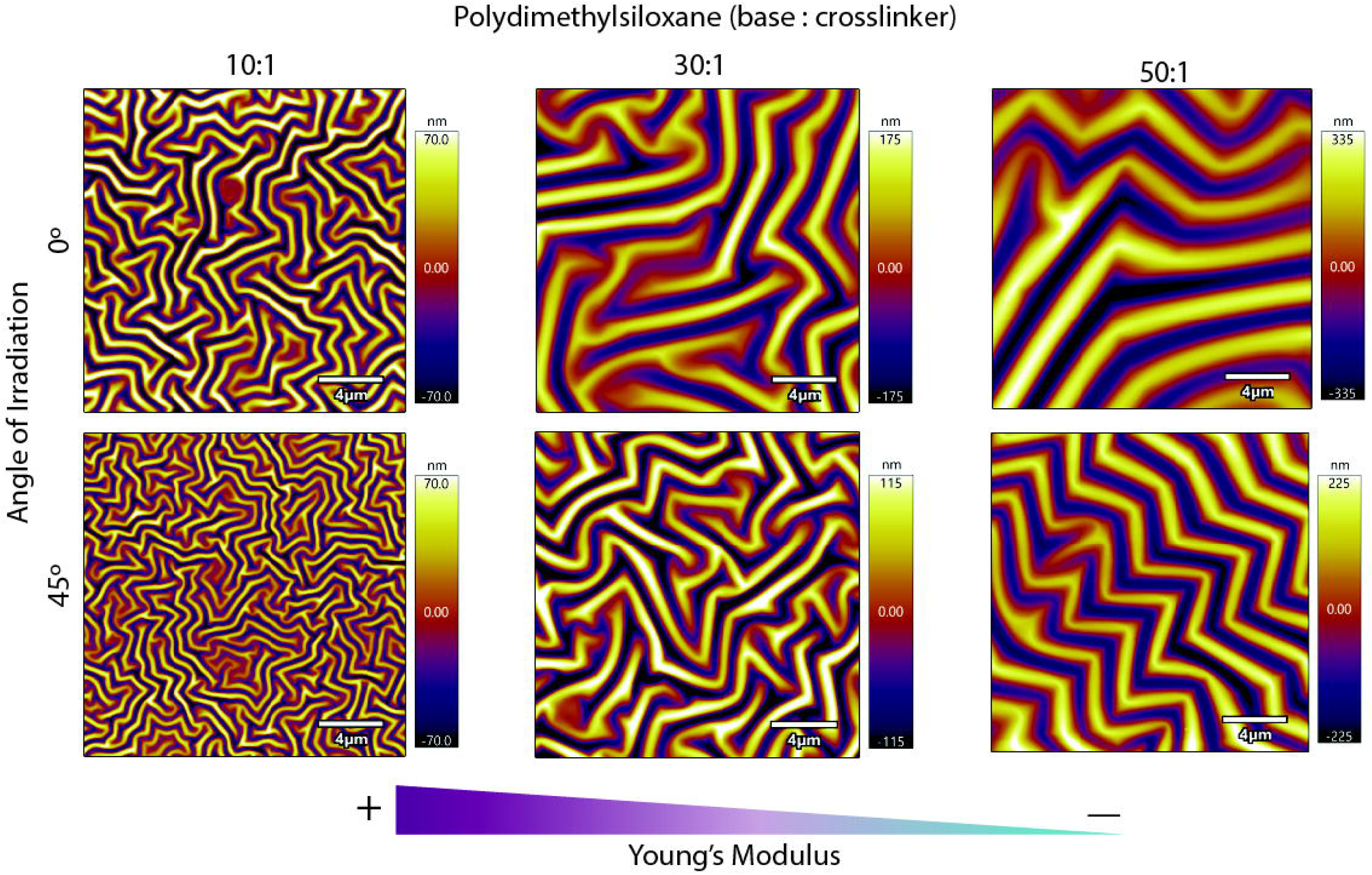
Wrinkle morphology of argon-irradiated PDMS as a function of the material stiffness and the angle of irradiation. The AFM images reveal that for the same angle of irradiation, the amplitude and wavelength of the wrinkles get larger as the Young modulus decreases from about 1 to 0.01 MPa. For a fixed crosslinked ratio, increasing the angle of irradiation reduces the wrinkle’s dimensions.

We performed an analysis of the power spectral density in reciprocal space to correlate the wavelength and height of the wrinkles with the nanofabrication parameters, the polymer Young’s modulus, and the resulting surface free energy (**Figure 2a**). The power spectral density describes how a signal’s power is distributed in frequency and decomposes the spectral content of the topography into contributions from different spatial frequencies or wavevectors in the form of peaks^27,28^. By extracting the area under the curve and the wavelength value at the center of each peak for each of the given PSD, we found that the root-mean-square (RMS) height and the frequency of the wrinkles increased monotonically from stiff to soft PDMS (**Figure 2b, c**). We also observed that the same wrinkled pattern could be obtained on substrates with different Young’s modules by manipulating the angle of irradiation. For example, the wrinkles obtained in PDMS with a 50:1 ratio and irradiated at an angle of 45° possess approximately the same wavelength and amplitude than the wrinkles obtained in 30:1 at 0°, despite the materials differs in stiffness by almost one order of magnitude (**Figure 2a-c**). The spectrum overlapping between these two substrates in the PSD plot further supports their morphological similarity (**Figure 2a**). The Young modulus of the PDMS increased after argon irradiation when compared to the untreated substrates across all the experimental samples, but it did not change significantly with the angle of irradiation for the same crosslinked ratio (**Figure 2c & Table 1**). In other words, for a given crosslinked density, the wrinkle size had little influence on the resulting PDMS stiffness, which is consistent with observations made by other authors^26^. Moreover, as the top layer of the PDMS crosslinked forming a silica-like thin film, we also expect the PDMS’s viscosity to decrease. **Table 1** gives a complete description of the irradiation parameters, wrinkle’s dimensions, and Young’s modulus for the different experimental samples. In our tests, the Young modulus was determined by using glass beads of 8 and 9.5 µm in radius. However, this method was not suitable for the measurement of Young’s modulus in untreated PDMS crosslinked at ratios of 30:1 and 50:1, even after submerging the sample in aqueous solutions, likely due to the stickiness and viscosity of the pristine polymer. Hence, the stiffness reported in **Table 1** for untreated PDMS at 30:1 and 50:1 ratio corresponds to values available in the literature^29^.

**Table 1.** Irradiation parameters used to produce wrinkle structures in PDMS with Young’s modulus ranging from about 0.01 to 1 MPa. The root-mean-square height, amplitude, and wavelength were extracted from the surface power spectral density. Values correspond to the mean ± standard deviation — ^□^ Young’s modulus reported by Ochsner and coworkers^29^.

| Wrinkled PDMS (Base: crosslinker) | Angle of Irradiation | RMS Height (nm) | Amplitude (nm) | Wavelength (μm) | Young's Mod. (MPa) |
| --- | --- | --- | --- | --- | --- |
| **10:1** | Untreated | - | - | - | 1.3±0.14 |
| | 0° | 40 ± 0.2 | 81 ± 0.3 | 0.83 ± 0.03 | 1.8±0.17 |
| | 45° | 33 ± 1.3 | 66 ± 2.3 | 0.60 ± 0.04 | 1.7±0.12 |
| **30:1** | Untreated | - | - | - | 0.1□ |
| | 0° | 77 ± 2.4 | 154 ± 4.7 | 1.97 ± 0.17 | 0.20±0.02 |
| | 45° | 68 ± 6.0 | 136 ± 12 | 1.34 ± 0.16 | 0.21±0.01 |
| **50:1** | Untreated | - | - | - | 0.01□ |
| | 0° | 173 ± 16 | 346 ± 33 | 3.10±0.18 | 0.07±0.002 |
| | 45° | 109 ± 2.7 | 218 ± 5.4 | 2.28±0.37 | 0.05±0.005 |

**Figure 2.**
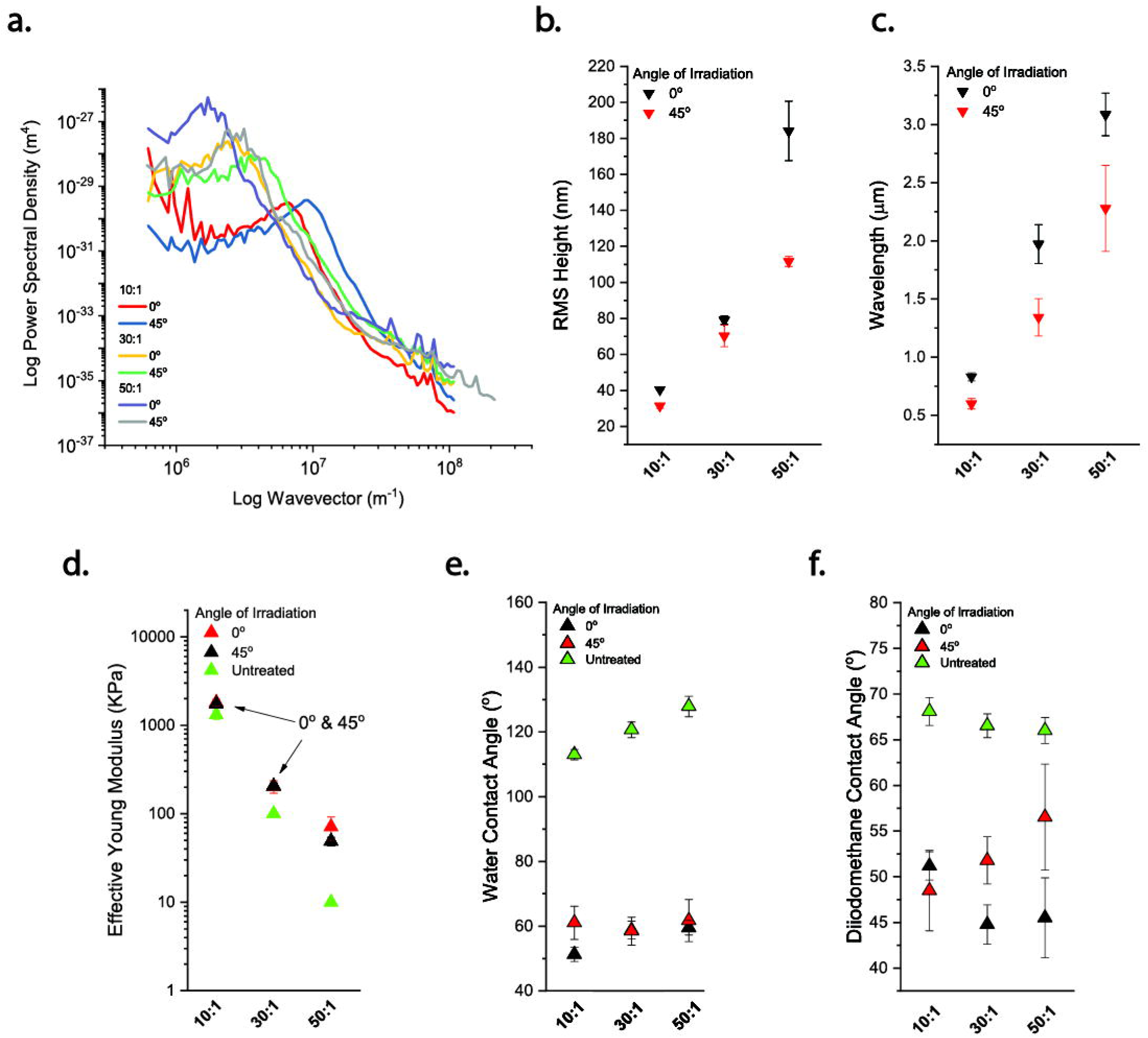
Argon-irradiation results in the formation of winkle structures in PDMS with the same wetting properties and comparable surface free energy but different Young’s moduli. **(a)** power spectral density (PSD) of wrinkled PDMS as a function of the Young modulus and angle of irradiation **(b)** Root-mean-square (RMS) height and **(d)** wavelength extracted from the PSD in (a). The RMS height and wavelength are larger when the stiffness of the PDMS decreases from about 1 MPa (10:1) to 0.01 MPa (50:1). **(c)** Effective Young’s modulus for different PDMS substrates. Notice that the changes in Young’s moduli as a function of the angle of irradiation are negligible. **(e)** The water contact angle of untreated and argon-irradiated PDMS. The wetting properties for the irradiated PDMS are analogous across all the experimental conditions. **(f)** The contact angle for diiodomethane (non-polar liquid), shows dependency on the material stiffness and angle of irradiation as the material elasticity decreases.

The PDMS substrates also became more hydrophilic after argon irradiation as the static water contact angle (WCA) decreased from about 110° to angles lower than 60° for all the irradiation conditions (**Figure 2e**), probably due to the higher oxygen content at the PDMS top layer that accompanied the oxidation of the polymer upon argon bombardment^26^. The inclusion of oxygen into the structure of the PDMS has been shown to add polarity to the polymer, thus reducing its hydrophobicity^30^ (**Figure 2d**). The WCA did not change significantly among samples with different stiffness and wrinkle sizes, except for the PDMS crosslinked at a ratio of 10:1 and irradiated at 0°, which had the lowest contact angle and was significantly different from the other irradiated samples (**Figure 2e**). These results indicated that argon irradiation of PDMS resulted in wrinkled structures with a variable size, but with the same stiffness and water wetting properties, despite the differences in surface stiffness and roughness. We also noticed that among the unmodified samples, the hydrophobicity raised monotonically as the ratio of base to crosslinker increased, and this difference was significant (*p*<0.05), contrary to the observations from other authors^31^. We further characterize the surface free energy of the irradiated PDMS by determining the contact angle of diiodomethane and solving for the surface energy according to the method developed by Owens and Wendt^24^, and recently verified by Hejda and coworkers^32^. This strategy uses a pair of non-polar (diiodomethane) and polar (water) liquids to get an estimation of the surface free energy. We found that the contact angle of the diiodomethane was dependent on the angle of irradiation and crosslinking ratio, which was probably associated with variations on the degree of oxidation of the PDMS^26^ (**Figure 2f**). However, we found that substrates with different stiffness and close nanoscale topography exhibited approximately the same surface energy (**Table 2**). For example, the PDMS crosslinked at 10:1 and irradiated at 45° had nearly the same surface free energy than the PDMS crosslinked at 30:1 and irradiated 0° and 45°. In general, the surface free energy varied between 42 to 51 mJ/m^2^ across all the wrinkled substrates. We did not find a solution for the surface free energy in the case of the untreated PDMS using the method developed by Owens and Wendt ^24^. Overall, we obtained substrates with variable substrate stiffness and with approximately the same interfacial energy by tuning the irradiation parameters.

**Table 2.** Components of surface energy for untreated and wrinkled poly(dimethylsiloxane) (PDMS) for different angles of irradiation and material stiffness. Values for the contact angle correspond to the mean ± SD. The surface energy was obtained according to the method developed by Owens and Wendt^24^. *The surface energy and its components correspond to mean values.

| Wrinkled<br>PDMS<br>(Base:<br>crosslinker) | Angle of<br>Irradiation | Contact Angle |  | *Surface energy [mJ/m<sup>2</sup>] |  |  |
| --- | --- | --- | --- | --- | --- | --- |
| | | Water | Diiodomethane | $\gamma_s^d$ | $\gamma_s^h$ | $\gamma_s$ |
| **10:1** | Untreated | 113 ± 1.6 | 68 ± 1.5 | - | - | - |
| | 0° | 51 ± 2.2 | 51 ± 1.5 | 26 | 25 | 51 |
| | 45° | 61 ± 5.1 | 48 ± 4.4 | 29 | 16 | 45 |
| **30:1** | Untreated | 121 ± 2.4 | 67 ± 1.3 | - | - | - |
| | 0° | 59 ± 2.7 | 45 ± 2.2 | 30 | 17 | 47 |
| | 45° | 58 ± 4.3 | 52 ± 2.6 | 26 | 20 | 46 |
| **50:1** | Untreated | 128 ± 3.1 | 66 ± 1.4 | - | - | - |
| | 0° | 60 ± 2.2 | 46 ± 4.4 | 30 | 17 | 47 |
| | 45° | 62 ± 6.5 | 57 ± 5.8 | 24 | 18 | 42 |

We started our analysis on bacterial adhesion and biofilm formation by studying the effect of substrate stiffness and wrinkle size on the ability of *E. coli* cells to colonize the different PDMS substrates. To that end, untreated and wrinkled PDMS samples were incubated with a suspension of exponential-phase *E. coli* cells at 30 °C for 24h in minimum media (M63). At the end of the incubation period, samples were gently rinsed to remove the loosely attached cells, and subsequently, cells were either imaged or stained with Crystal Violet to determine biofilm mass accumulation. We found that the wrinkled topography reduced significant biofilm mass regardless of the PDMS stiffness, except for the softest wrinkled PDMS at 50:1 ratio (**Figure 3a**). We also observed that wrinkles with a wavelength of about 1 µm, which formed in PDMS at 10:1 ratio and 0° angle of irradiation, yielded a 2-fold biofilm reduction compared to the untreated sample at the same cross-linked ratio (**Figure 3a**). Also, biofilm reduction in this sample was significantly lower than the amount of biofilm formed in the softest wrinkled surface (50:1) but comparable to that of wrinkled PDMS at a 30:1 ratio. Laser confocal fluorescent images revealed that the distribution of bacteria became increasingly uneven on the wrinkled substrates, probably contributing to the decrease in biofilm mass measured by Crystal Violet (**Figure 3b**). Optical inspection showed that *E. coli* attached predominantly to the valleys and in direct contact with the material (**Figure 3c**). Few cells were detected on the wrinkle’s crest and sidewalls, even for the softest wrinkled PDMS that had the longest wavelength and the most extensive surface area for cell attachment (**Figure 3c**). Similarly, for wrinkles approaching the size of *E. coli* in the wrinkled PDMS at 10:1 ratio, the cells settled inside the wrinkle’s trough despite the cost of bending or aberrant cell shapes induced by the microtopography (**Figure 3d**).

**Figure 3.**
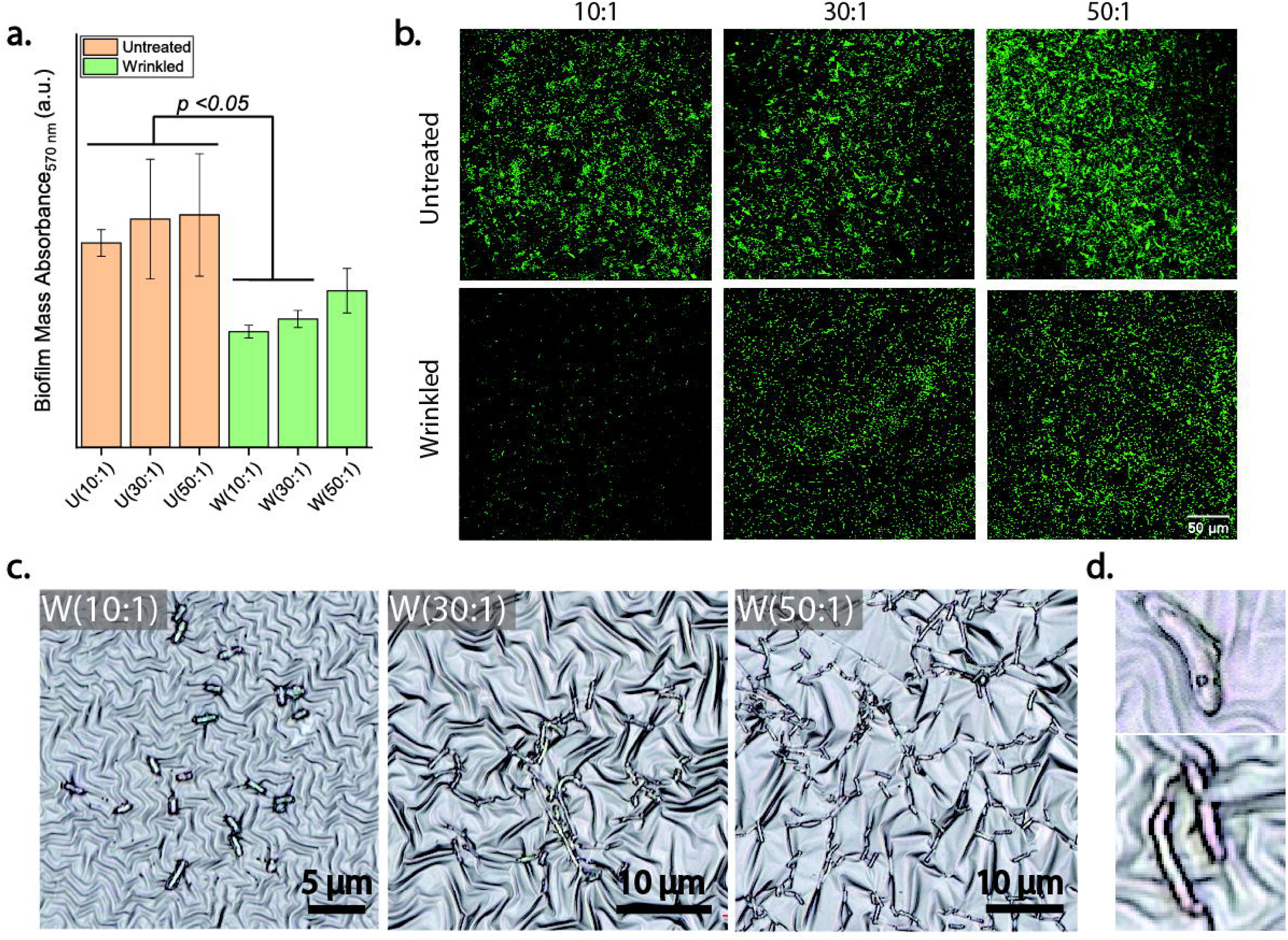
Surface topography reduces *E. coli* adhesion and biofilm formation compared to the untreated PDMS. **(a)** Relative biofilm mass assessed by Crystal Violet staining. Crosslinking ratios of 10:1, 30:1 and 50:1 correspond to Young’s modulus of about 2 MPa, 0.2MPa, and 0.02MPa, respectively, for wrinkled PDMS. Untreated (U) and wrinkled (W) PDMS was cultured with *E. coli* bacterial suspensions for 24 h at 30°C. In the bar plot: biofilm mass accumulation is relative to the untreated sample, a.u. indicates arbitrary units, and *p<* 0.05 denotes significant differences between samples. **(b)** Representative fluorescent images of *E. coli* constitutively expressing the Green Fluorescence Protein (GFP). **(c)** Representative optical images of *E. coli* respect to the wrinkled topography; **(d)** optical images of wrinkled PDMS crosslinked at 10:1 ratio, showing the distortion of the bacterial cell shape imposed by the sharp turns and tortuosity of the topography.

The differences in bacterial adhesion observed on the softest wrinkled PDMS at 50:1 ratio might stem from the large wrinkled size in this sample compared to that on the 10:1 and 30:1 irradiated substrate, but not from differences in material stiffness. If this was the case, we would expect to see a measurable decrease in biofilm mass as the wrinkle size decreases. To test this possibility, we cultured *E. coli* cells on 50:1 and 30:1 PDMS that were irradiated at angles of 0° and 45° to yield wrinkles with a shorter wavelength (see **Figure 1**). As previously detailed, the 50:1 PDMS irradiated at 45° reproduces the same topography than on the 30:1 substrate irradiated at 0°, despite the differences in the material stiffness. Also, the wrinkle size does affect the Young modulus in the wrinkled PDMS for the same crosslinking ratio (**Figure 1**). We found that the biofilm mass formation diminished significantly in both cases by making the wrinkles shorter, supporting our initial observations (**Figure 4a**). This also suggests that even for very soft PDMS, biofilm mass can be mitigated by tuning the surface topography alone. To further support this notion, we compared the wrinkled structures formed at a 10:1 ratio, which yielded the lowest biofilm mass (**Figure 3a, b**), with those formed in the 30:1 PDMS irradiated at angles of 45° and 75°. Those wrinkled substrates were selected because they have a comparable surface free energy (see **Table 2**); thus, ruling out the influence of this parameter on adhesion and biofilm initiation of *E. coli* on PDMS with very dissimilar stiffness (≈ 2 vs. 0.2 MPa). As aforementioned, the wetting properties for the untreated PDMS was significantly different across the crosslinked densities used here. Using this experimental setup, we found that biofilm mass formation was not significant between the PDMS with Young’s modulus of 1.8 and 0.2 MPa, which corresponds to crosslinked ratios of 10:1 and 30:1, respectively. Also, under these conditions, as shown in **Figure 3**, *E. coli* can still make direct contact with the underlying topography.

**Figure 4.**
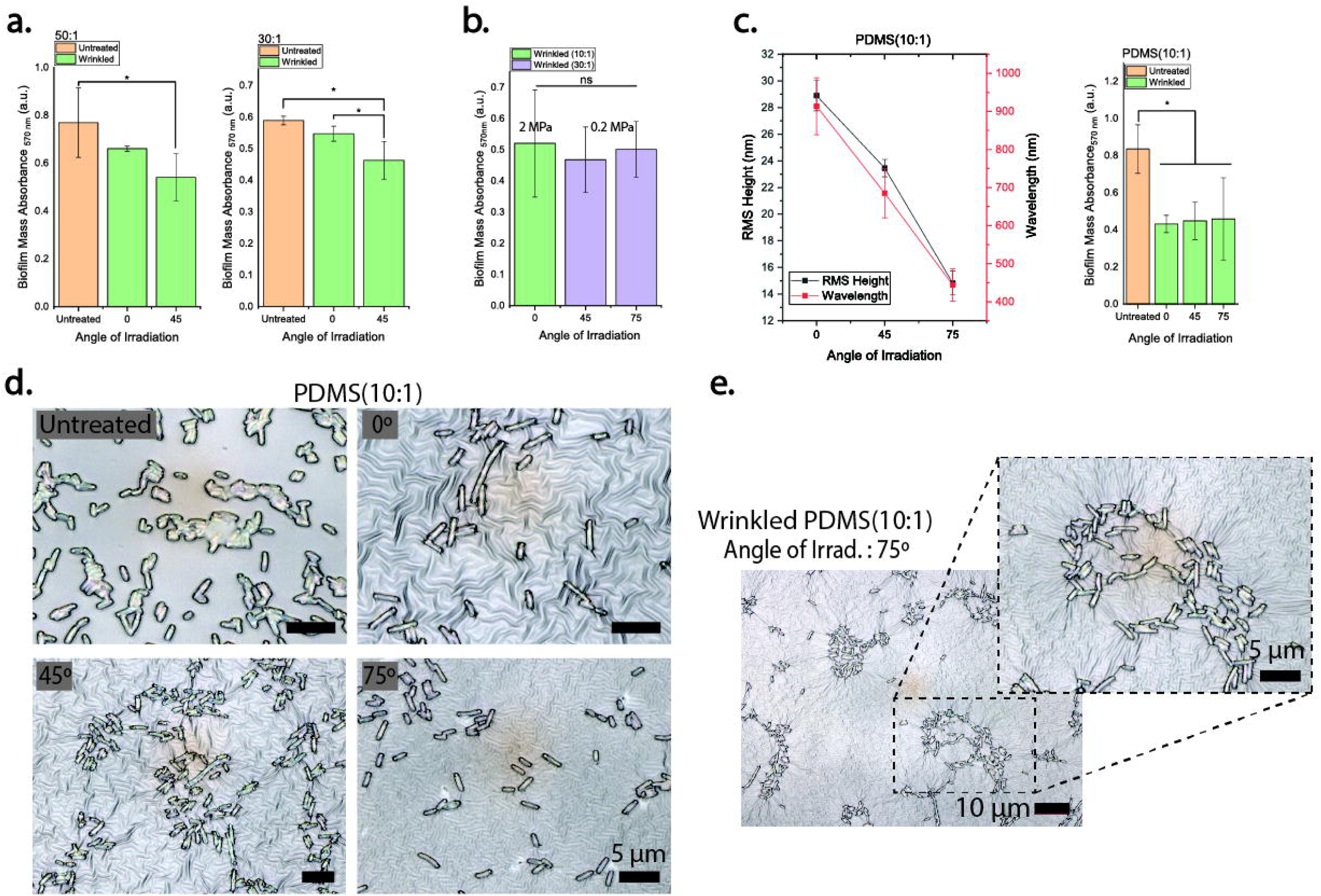
Surface topography reduces *E. coli* adhesion and biofilm formation on both stiff and very compliant PDMS. **(a)** Crystal Violet staining of biofilms formed on very compliant PDMS with decreasing wrinkled dimensions. Notice that biofilm mass accumulation was reduced in both 30:1 and 50:1 by decreasing the wrinkle size. **(b)** The data do not show any significant differences between the wrinkled PDMS at 30:1 and 10:1 ratio irradiated at oblique angles. **(c)** For all the bar plots: biofilm mass accumulation is relative to the untreated sample, \**p<* 0.05, a.u. indicates arbitrary units, and n.s. denotes no significant differences between samples. **(d)** Optical images of *E. coli* seeded on a substrate with stiffness of about 2 MPa with decreasing wrinkle’s dimensions. **(e)** *E. coli* accumulation in topographical defects found in the 10:1 PDMS irradiated at 75°. The inset is an enlargement of *E. coli* settling down in these regions.

Lastly, we investigated if decreasing the wrinkles to sizes much smaller than the *E. coli* dimensions would cause an opposite effect on bacterial adhesion, that is, increase biofilm mass as the surface approaches to a flat surface. To that end, 10:1 PDMS was irradiated at angles of 0°, 45°, and 75° to yield wrinkles with RMS heigh and wavelength at the submicron scale (**Figure 4b**). Biofilm mass formation was quantified by Crystal Violet, following the same procedure as described previously. Interestingly, we found that in all the cases, the nanotopography significantly reduced biofilm mass formation by two-fold when compared to the untreated control, even when the wrinkle’s size approached 400 nm in wavelength (**Figure 4c**). However, this sample also showed the highest standard deviation in contrast to the other wrinkled structures (formed at 0° and 45°). Optical inspection revealed that while *E. coli* settled down mostly on topographical defects that were present on the 75°-wrinkled PDMS (**Figure 4e**). This data suggests that when the wrinkled pattern has dimensions smaller than *E. coli*, the patterned PDMS films may have the best performance on preventing biofilm mass accumulation.

## Discussion

Microbial contamination of biomaterial surfaces underpins the pathogenesis of device-associated infections and constitutes an adverse event that compromises patient safety and material functionality. Device-associated infections are impressively resilient to antibiotic treatment alone, and thus in most cases, their management requires biomaterial removal, which increases the risk of patient morbidity and mortality and excess costs to the healthcare system^1^. Device-associated infections often precede other life-threating complications, including bacteremia, catheter-related bloodstream and urinary tract infections, and ventilator-associated pneumonia. Bacterial adhesion to the biomaterial surface is the first event leading biofilm formation; therefore, understanding the molecular and physical interactions that underlie bacteria-surface interactions are pivotal to design strategies that can prevent the formation of biofilms on the biomaterial surfaces^3^. In this context, material stiffness has been recently suggested to influence microbial adhesion and their biofilms in a variety of polymers and hydrogels. However, whether very compliant substrates are deleterious or otherwise advantageous to bacterial cell adhesion are still debatable, and no consensus exists about the role of material stiffness on biofilm initiation. One reason for these disparities is possibly due to the influence of confounding variables like viscosity, porosity, and hydrophobicity, which arise when making polymeric preparations with a wide range of composition-stiffness combinations, masking the effect that substrate elasticity may have in bacterial physiology.

To shed light on this matter, we irradiated PDMS with singly-charged argon ions to yield substrates with approximately the same surface free energy but variable substrate stiffness. This process also resulted in the formation of micron and submicron wavy structures (wrinkles), whose dimensions were tuned by controlling the irradiation parameters (e.g., angle of irradiation). We have recently demonstrated that ion-beam irradiation can also be leveraged to induce the growth of high aspect ratio nanostructures in a hydrogel^33^. Interestingly, the wrinkle’s dimensions had no effect in the resulting PDMS stiffness for a fixed crosslinked ratio. Moreover, by tuning the angle of irradiation, the same wrinkle morphology was obtained in PDMS with elasticities differing by one-order of magnitude. By making PDMS preparations at different crosslinked ratios but analogous surface free energy, we found that biofilm mass accumulation was distinctly affected by the surface topography and not by the material stiffness. Specifically, by designing wrinkled structures with sizes comparable to the bacterium, biofilm mass accumulation was not statistically significantly different between PDMS with Young’s modulus of 0.02, 0.2, and 2 MPa, all of them having a surface free energy in the range of 4.2-5.1 mJ/m^-2^.

Although our findings contrast with reports suggesting that material stiffness directly impacts cell attachment and biofilm formation^20,31^, those can be reconciled if one considers, for example, the role that other surface properties have on microbial adhesion including the interfacial energy, as the polymer crosslinking density varies. Indeed, we found that the wetting properties for the untreated PDMS were (i) significantly different in preparations with elastic modulus differing in one and two orders of magnitude, and (ii) that material hydrophobicity increased monotonically as the compliance of the PDMS augmented. Even though we did not measure material viscosity, our results are in agreement with those reported by Straub and coworkers^21^. These authors have suggested that varying the crosslinking ratio of polymers such as PDMS, leads to preparations with high hysteresis and large interfacial stickiness, which may enhance bacterial retention. The same authors have further suggested that the high bacterial adhesion to very compliant PDMS (e.g., preparations of the base to crosslinker agent of 50:1) likely stems from strong hydrophobic interactions between the bacterium and the polymer.

Despite the sharp turns and tortuosity of the wrinkled topography in the irradiated PDMS, we also found that *E. coli* adapted its morphology to fit inside the wrinkle’s trough when the feature dimensions were larger or comparable to those of the bacterium. It has been demonstrated that *E. coli* can plastically deform and adapt its growing morphology when confined in microcavities^34^, exposed to bending forces^35^, or be challenged by mechanical compression^36^, and recover its straight, native rod-like morphology when the external force is removed. Our results agree with other reports showing that bacterial cells selectively colonize receding topographical features at the micron and submicron scale^37,38^. The observed preferential settlement of *E. coli* into the wrinkle’s troughs could also be due to the removal of *E. coli* from the wrinkle’s crests during sample rinsing, or due to unfavorable attachment on those curved areas. In this regard, Chang and coworkers^11^ observed that surface motility of *Pseudomonas aeruginosa* was affected by the radius of curvature of the micrometer-scale hemispherical topographies and suggested that bacterial adhesion to those areas was energetically unfavorable since it required cell bending. Interestingly, we observed that when the wrinkles became much smaller than the bacterial dimensions and at the submicron scale, the anti-biofouling properties of the material were comparable to those features with about 1 µm in wavelength. However, a closer examination showed those similarities probably arose from topographical defects in the sample after irradiation, allowing *E. coli* to establish in those areas. In agreement with our observations, other authors had reported that when the pattern size was shorter than the bacteria size (four-way grids of 0.5 µm in size), the anti-biofouling capacity of the films increased significantly^39^. It would be interesting to determine if the geometry of the topography (e.g., curvature) at the submicron scale might contribute to the observed effect.

In summary, we were able to fabricate PDMS with a stiffness that spanned several orders of magnitude, i.e., from few kPa to several MPa, and analogous surface free energy (ranging from 42 to 51 mJ/m^2^). Low-energy ion beam irradiation also resulted in the crosslinking of the polymer’s top layer, producing wrinkles whose amplitude and wavelength were tuned by controlling the angle of irradiation. Our results suggest that topographical patterns impart anti-biofouling properties to a surface against *E. coli* that are independent of substrate stiffness. We also demonstrate that topography can limit bacterial surface attachment and biofilm formation even in very compliant PDMS (Young’s modulus of 0.02 and 0.2 MPa). At first sight, our findings contrast with other reports suggesting a direct role of substrate stiffness on bacterial adhesion and biofilm initiation. However, those findings can be reconciled if one considers that the adherence of biofilms to surfaces is also governed by hydrophobic interactions, protein adhesion, van der Waals forces, electrostatic, and steric interactions. Because *E. coli* is a significant source of medical-device acquired infections, we believe our findings are a cost-effective and simple way to reduce bacterial attachment before biofilms can be formed, even in very compliant and hydrophobic polymers.

## Acknowledgments

Material characterization was carried out in part in the Materials Research Laboratory Central Research Facilities at the University of Illinois at Urbana-Champaign. Laser scanning confocal fluorescent microscopy was conducted at the Microscopy Suite of the Beckman Institute for Advanced Science and Technology at the University of Illinois at Urbana-Champaign (UIUC-BI-MS).

